# Delineating transcriptomic signatures of in vitro human skeletal muscle models in comparison to in vivo references

**DOI:** 10.1101/2025.02.28.640821

**Authors:** Margaux van Puyvelde, Eslam Essam Mohammed, Ángela Moreno Anguita, Jarne Bonroy, Alexandra Jansen, Atilgan Yilmaz

## Abstract

A pivotal question at the heart of stem cell research is how faithful cellular models recapitulate the biology of human tissues. Skeletal muscle is the largest tissue in the human body and has been extensively modelled by various in vitro systems. Here, we sought to delineate the state-of-the-art of in vitro human skeletal muscle models by performing a large-scale analysis of transcriptome datasets, covering more than 400 samples across 39 studies, including bulk and single cell RNA sequencing of 2D and 3D models and their in vivo counterparts. Our analyses highlight common discrepancies between a wide range of cellular models and human skeletal muscle with respect to their myogenic identity, transcription factors, epigenetic complexes, metabolic processes and signaling pathways. Our analyses reveal cellular processes that can be modulated to improve the in vitro models of skeletal muscle, while also paving the way for similar studies for other cell types.

## Introduction

The use of reliable in vitro models of human tissues is essential to study development, disease phenotypes, regeneration and putative therapeutic interventions. Methods of model generation vary from the differentiation of pluripotent stem cells to transdifferentiation of somatic cells or the use of primary isolated cell types, which can be immortalized^1–7^. Despite the great technical advantages they provide and a vast number of insights they have given into human development and disorders, all of these cellular models have their own challenges. Primary cells quickly lose many of their in vivo characteristics and after a prolonged period of culturing, will enter senescence. Immortalizing the primary cells by overexpression of *TERT*, *CDK4* and *SV40* extends the lifespan of the cells but also alters their expression patterns^8^. While differentiation protocols to any cell type from human pluripotent stem cells are often long and expensive, transdifferentiation from somatic cells typically exhibits low efficiency levels with partial retention of the epigenetic profile of the source cells. Additionally, the cells obtained through differentiation and transdifferentiation methods have been described to resemble more of a fetal-like identity^9^. Although the improvements in stem cell-based in vitro models depend on a thorough assessment of cellular identities of these models at the molecular level, a systematic characterization of their differences in comparison to bona fide human cell types remains to be done.

Skeletal muscle is the largest tissue, encompassing about 40 percent of the human body mass^10,11^. Not only does it provide the mechanism behind movement, but it also plays an essential role in metabolism and regulates immune system functions. Skeletal muscle is susceptible to a plethora of genetic and metabolic disorders, while also undergoing wasting in cancer and upon aging, making this tissue one of the prime targets of regenerative medicine. Thus, having 2D and 3D models that can faithfully recapitulate human muscle is crucial to get insight into the development, diseases and regeneration of this tissue and to aid the identification of novel therapeutic interventions.

A growing body of research has been dedicated to the analysis of skeletal muscle models and biopsies through the lens of transcriptomics. While the majority of the efforts were focused on bulk RNA sequencing, a smaller and more recent pool of studies made use of single cell RNA (scRNA) sequencing. Transcriptome analysis has been instrumental in elucidating developmental trajectories and identifying disparities between healthy and diseased muscle tissue. Nevertheless, to date, skeletal muscle transcriptomic data has not been utilized to discern molecular differences between the in vitro models and bona fide skeletal muscle samples in a systematic way.

In this study, we bring together 39 bulk and single cell RNA sequencing studies covering over 400 samples from all types of in vitro skeletal muscle models and compare these to different stages of human adult and fetal muscle biopsies^12–48^. We show failure of expression in several myogenic factors, aberrant transcription factor signatures, epigenetic memory retention and major differences in fatty acid metabolism and membrane transporter expression patterns in different in vitro models. Interestingly, the integration of 6 single cell RNA sequencing datasets covering 2D and 3D human pluripotent stem cell-derived skeletal muscle progenitors, and their adult, fetal and embryonic counterparts revealed different degrees of quiescence in *PAX7^+^* cell populations derived from different methodologies. Furthermore, integration of single cell transcriptome datasets also uncovers a potential function for BRCA1-BRCA2-containing Complex in proliferating developmental myogenic progenitors in human. Our analyses shed light on the common discrepancies between in vitro models and bona fide skeletal muscle cells across different cellular processes and provide a reference for future studies to improve the existing models.

## Results

### Large-scale analysis of bulk RNA sequencing samples reveals differences between in vitro models and bona fide skeletal muscle

To identify the differences between the in vitro models of human skeletal muscle cells and their in vivo counterparts, we assembled a comprehensive dataset of more than 400 samples from 34 studies using bulk RNA sequencing and 5 studies using single cell or single nucleus RNA sequencing. This curated dataset covers a wide range of in vitro skeletal muscle cell models together with their source materials as well as in vivo human skeletal muscle samples derived from embryonic, fetal and adult stages of life. Thus, our analyses included human pluripotent stem cells (hPSC), hPSC-derived myogenic progenitors, 2D and 3D hPSC-derived myotube cultures, fibroblasts, fibroblast-derived transdifferentiated myoblasts and myotubes, adult tissue-derived myogenic progenitors, fetal and embryonic myogenic progenitors, adult isolated myofibers, heterogeneous adult and fetal biopsies, immortalized myogenic cell lines and primary cultures of human muscle cells (Figure 1a).

**Figure 1.**
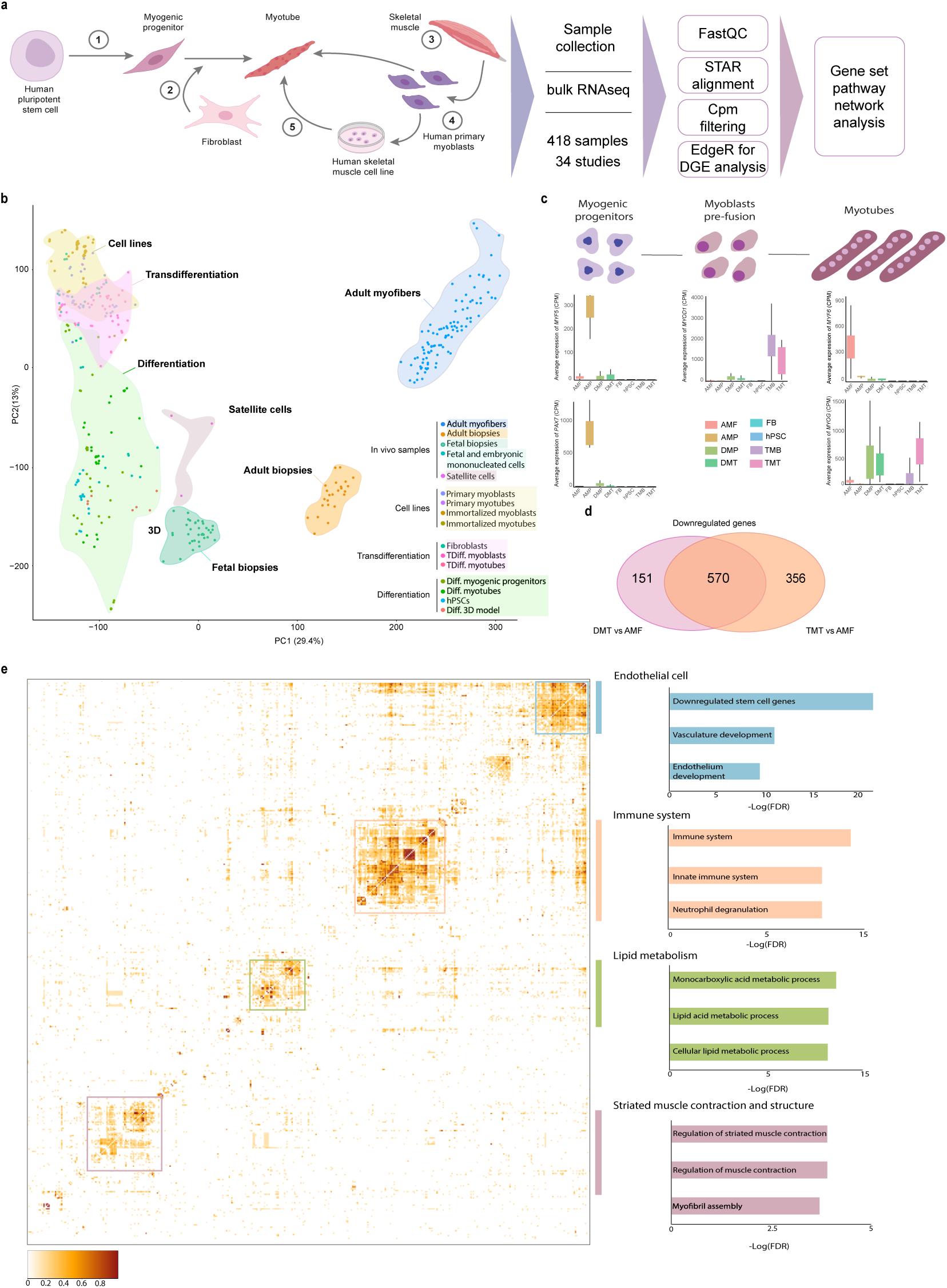
Transcriptome-wide comparison and myogenic profiles of the integrated in vitro and in vivo skeletal muscle samples. **a)** Schematic overview of sample collection and analysis pipeline. **b)** Principal component analysis (PCA) for all collected samples included in the study. **c)** Average counts per million (CPM) values of the myogenic regulatory factors (MRFs) for their respective stages depicted in the illustration for hPSC-derived differentiated myogenic progenitors (DMP) and myotubes (DMT), transdifferentiated myoblasts (TMB) and myotubes (TMT), fibroblasts (FB), human pluripotent stem cells (hPSCs), adult myogenic progenitors (AMP) and adult isolated myofibers (AMF). **d)** Venn diagram showing the overlap in significantly downregulated genes (FDR<0.05) between differentiated myotubes and transdifferentiated myotubes as compared to adult isolated myofibers. **e)** Heatmap demonstrating predicted protein-protein interaction network scores for the commonly downregulated genes in Fig. 1d. Predicted interaction values represent a confidentiality score between 0 and 1, as depicted in the scale bar. Strong interaction clusters are analyzed through Gene Set Enrichment Analysis (GSEA) and are summarized in the bar plots on the right of the heatmap. For each gene ontology term with significance cut-off of *P* < 0.05, the negative standard logarithm of the adjusted *P* value (-log(FDR)) is plotted.

A principal component analysis (PCA) of all bulk RNA sequencing samples demonstrated a clear divide between the in vitro models and the in vivo reference samples, with the adult samples separating from the rest of the samples and the in vitro samples positioning closer to fetal biopsies (Figure 1b). Next, we were interested in investigating the expression levels of the myogenic regulatory factors (MRFs), which are a set of transcription factors that have master regulatory roles during the highly orchestrated process of myogenesis. Our dataset covers different stages of myogenesis by profiling the myogenic progenitors, myoblasts and myotubes for the in vitro systems and the in vivo references. During the myogenic progenitor stage, there is clear upregulation of *MYF5* and *PAX7* in the hPSC-derived differentiation model and the adult myogenic progenitors, however, expression of these myogenic markers is virtually absent in the transdifferentiation model. *MYOD1* marks the transition of myogenic progenitors towards myoblasts and primes the progenitors for fusion. As expected, this MRF is highly upregulated in the transdifferentiation model owing to the use of the inducible overexpression of *MYOD1* as a common methodology to reprogram fibroblasts into the myogenic lineage. *MYOG* expression is present in the mature stage of both in vitro models and the in vivo references, whereas *MYF6* failed to be expressed specifically in the transdifferentiation model (Figure 1c).

To get a first insight into the main differences between hPSC-derived differentiated and fibroblast-derived transdifferentiated myotubes in comparison to isolated adult human myofibers, we examined the overlap in downregulated differentially expressed genes (DEGs) between the two models. Since myotube cultures are typically heterogenous, we focused on downregulated DEGs with strict expression criteria (CPM < 1 in vitro) to identify the genes that are virtually absent in the in vitro models as opposed to their expression in vivo. In addition, our dataset includes several studies with different in vitro protocols, which further consolidates the confidence in the commonly downregulated genes across these protocols. In total, hPSC-derived differentiated and fibroblast-derived transdifferentiated myotube cultures showed an overlap of 570 DEGs, highlighting a striking overlap of more than 60% of their total number of DEGs (Figure 1d). To further investigate this overlapping gene group, we analyzed their predicted protein-protein interactions using STRING (Figure 1e, left). This analysis revealed four groups of genes that are predicted to be interacting highly as an interconnected group. To get insight into the potential functions of the genes in these interaction nodes, we performed Gene Set Enrichment Analysis (GSEA). Enriched gene ontology terms for these four major gene groups suggested a function for these genes in striated muscle contraction and structure, lipid metabolism and surprisingly also endothelial cells and the immune system (Figure 1e, right).

To rule out the possibility of a major contamination of immune and endothelial cells in the isolated adult myofibers, we analyzed a recent single nucleus RNA sequencing dataset of a complete adult muscle biopsy for the expression of the genes predicted to be related to these cell types^49^ (Supplementary Figure 1a). 25% percent of the genes enriched in the gene ontology terms related to these cell types were also simultaneously expressed in the myofibers. 10 of these genes were robustly expressed at high levels in the myofiber-associated nuclei (Supplementary Figure 1b-k), suggesting previously uncharacterized functions for these genes within adult myofibers. 20 additional genes showed low to medium expression within myofiber-associated nuclei (Supplementary Figure 1l). In summary, a first global look across the MRFs and the genes that fail to be expressed in both in vitro systems show that in vitro models exhibit disparities compared to the in vivo references, although they resemble the fetal stages more. They also lack the expression of structural and lipid metabolism-related genes associated with adult skeletal muscle in human and a group of previously overlooked genes.

### Aberrant expression of transcription factors and epigenetic complexes in the in vitro models

Subsequently, we sought to investigate the expression of transcription factors and epigenetic complexes in the in vitro models in comparison to the in vivo references, as these molecules are the major drivers of cell fate changes. We thus filtered the DEG lists by a comprehensive list of human transcription factors and epigenetic factors derived from EpiFactors database. This analysis revealed two major transcription factor families as being differentially expressed in the in vitro models. Multiple members of the HOX family of transcription factors were consistently upregulated in both myoblast and myotube stages of hPSC-derived differentiation and fibroblast-derived transdifferentiation models in comparison to isolated adult human myofibers (Figure 2a). The same trend was also recapitulated in the immortalized cell line myotubes, in particular for the case of several members of the HOXB cluster (Supplementary Figure 2a). HOX genes have a well-described role in spatial patterning during development and control muscle diversity, therefore, they are likely to also regulate initial fate specification in vitro^31,50^. Additionally, we found significant differential expression of the members of Ankyrin Repeat and Death Domain Containing (ANKRD) transcription factor family, of which two have been described to play important roles in skeletal muscle, namely *ANK3* and *ANKRD2 (*Figure 2a)^51,52^.

**Figure 2.**
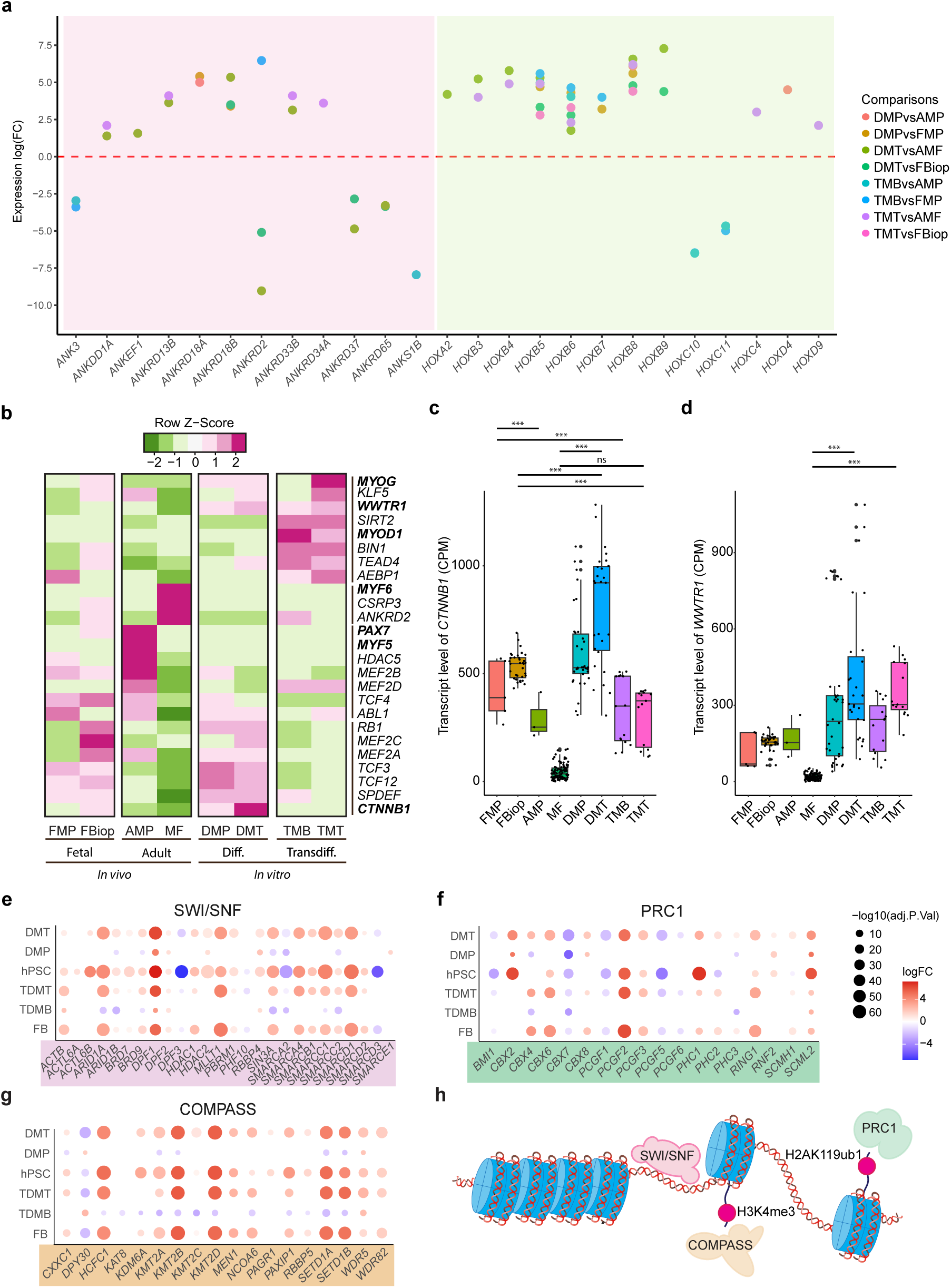
Analysis of expression of transcription factors and epigenetic complexes in the in vitro models. **a)** Dot plot showing the positive and negative standard logarithmic fold change of gene expression for differentially expressed genes across the indicated comparisons. Each unique comparison is color-coded, and genes are grouped on the x-axis based on the transcription factor families they belong to, with the Ankyrin Repeat and Death Domain Containing gene family being on the left and HOX family on the right. **b)** Heatmap demonstrating the average CPM values for transcription factors associated with the myogenic identity. Highlighted in bold are MRFs, a Hippo pathway effector, *WWTR1*, and the key component of WNT pathway, *CTNNB1*. **c-d)** Box plots showing expression levels of *CTNNB1* and *WWTR1,* across all categories. **e-g)** Dot plot demonstrating the expression of the active members of the SWI/SNF **(e)**, PRC1 **(f)** and COMPASS/MML **(g)** complexes across three stages of hPSC-derived differentiation and fibroblast-derived transdifferentiation models. Source material, which are hPSCs and fibroblasts, and in vitro generated myotube samples were compared to isolated adult human myofibers, whereas the mononucleated myogenic intermediates were compared to isolated mononucleated myogenic progenitors from adult biopsies. Dot colors indicate standard logarithmic fold change and dot size is determined by the negative standard logarithm of the adjusted *P* values for the respective fold change. **h)** Schematics illustrating the canonical chromatin function of the epigenetic complexes highlighted in Fig. 2e-g.

We then focused on the expression patterns of the transcription factors that are canonically associated with myogenic identity. In addition to the MRFs that we found to be differentially expressed in Figure 1c, the key downstream effector of Wnt pathway, *CTNNB1,* and the Hippo signaling co-factor, *WWTR1,* also showed aberrant expression patterns in the in vitro models (Figure 2b-d). *CTNNB1*, which encodes β-catenin, was expressed at high levels in hPSC-derived differentiation model, whereas it showed very low expression levels in adult myofibers, suggesting the possibility of a sustained, aberrant Wnt activity in the in vitro systems. Importantly, the small molecule, CHIR99021, which activates the Wnt pathway through GSK-3 inhibition, has been suggested to enhance the efficiency of transdifferentiation and is also commonly used in widely accepted hPSC-derived skeletal muscle differentiation protocols^2,53^. Similarly, *WWTR1* showed higher expression levels in both in vitro models as compared to fetal or adult references. Finally, we found that several active members of three major epigenetic complexes, SWI/SNF, PRC1 and COMPASS had altered expression patterns in the in vitro models as compared to their respective references throughout the different stages of myogenesis starting from their source material (Figure 2 e-h). Both SWI/SNF and PRC1 have been shown to play pivotal roles in myogenesis and cell differentiation^54,55^. Therefore, dysregulation of their active members might affect the efficiency of generation of in vitro skeletal muscle. In both hESC-derived differentiation and fibroblast-derived transdifferentiation models, several genes associated with SWI/SNF, PRC1 and COMPASS/MLL complexes retained their expression from the source stage, i.e. hPSCs and fibroblasts, throughout the in vitro myoblast and myotube stages, suggesting an aberrant retention of epigenetic memory (Supplementary Figure 2 b-d).

### Dysregulation of metabolic homeostasis and fiber type signatures of in vitro models

The different stages of myogenesis are supported by specific changes in the metabolism. This process, referred to as metabolic reprogramming, is also a major component of muscle differentiation by switching progenitors from a quiescent to an active state in the adult stem cell population^56^.

To get insight into potential metabolic differences between the in vitro models and the in vivo references, we first investigated global changes of expression in metabolism-related genes. Across all metabolism-related genes, transmembrane transporters were found as a significantly enriched group that was differentially expressed across all stages of in vitro models (Figure 3a-b). Interestingly, metabolism-related DEGs were also enriched within gene ontology terms related to folic acid metabolism. Indeed, both the folic acid receptor *FOLR1* and three other active members of the pathway, *RAC1*, *RHOA* and *ROCK1*, were significantly upregulated in hPSC-derived differentiation model, while they showed a more striking upregulation in the transdifferentiation model (Figure 3c). In addition, members of several phases of the lipid cycle, including fatty acid catabolism, long chain fatty acid and very-long-chain fatty acid synthesis and phosphatidyl choline synthesis, were downregulated at the terminally differentiated myotube stages of hPSC-derived differentiation and fibroblasts-derived transdifferentiation models in comparison to the isolated adult myofibers (Figure 3d-e and 3g-h). Conversely, expression of members of the cholesterol synthesis pathway were downregulated in both of the in vitro models of human multinucleated myotubes (Figure 3f and 3h). In accordance with these observations, expression of members of folic acid metabolism and lipid cycle was also upregulated in the myotubes derived from immortalized myogenic cell lines as compared to adult isolated myofibers, while the members of cholesterol synthesis pathway were downregulated (Supplementary figure 3a and 3c-f). Both the upregulation of the folic acid metabolism and lipid cycle might argue for a failure in metabolic reprogramming or can be caused by media conditions, suggesting a need for adjustments in the media formulations.

**Figure 3.**
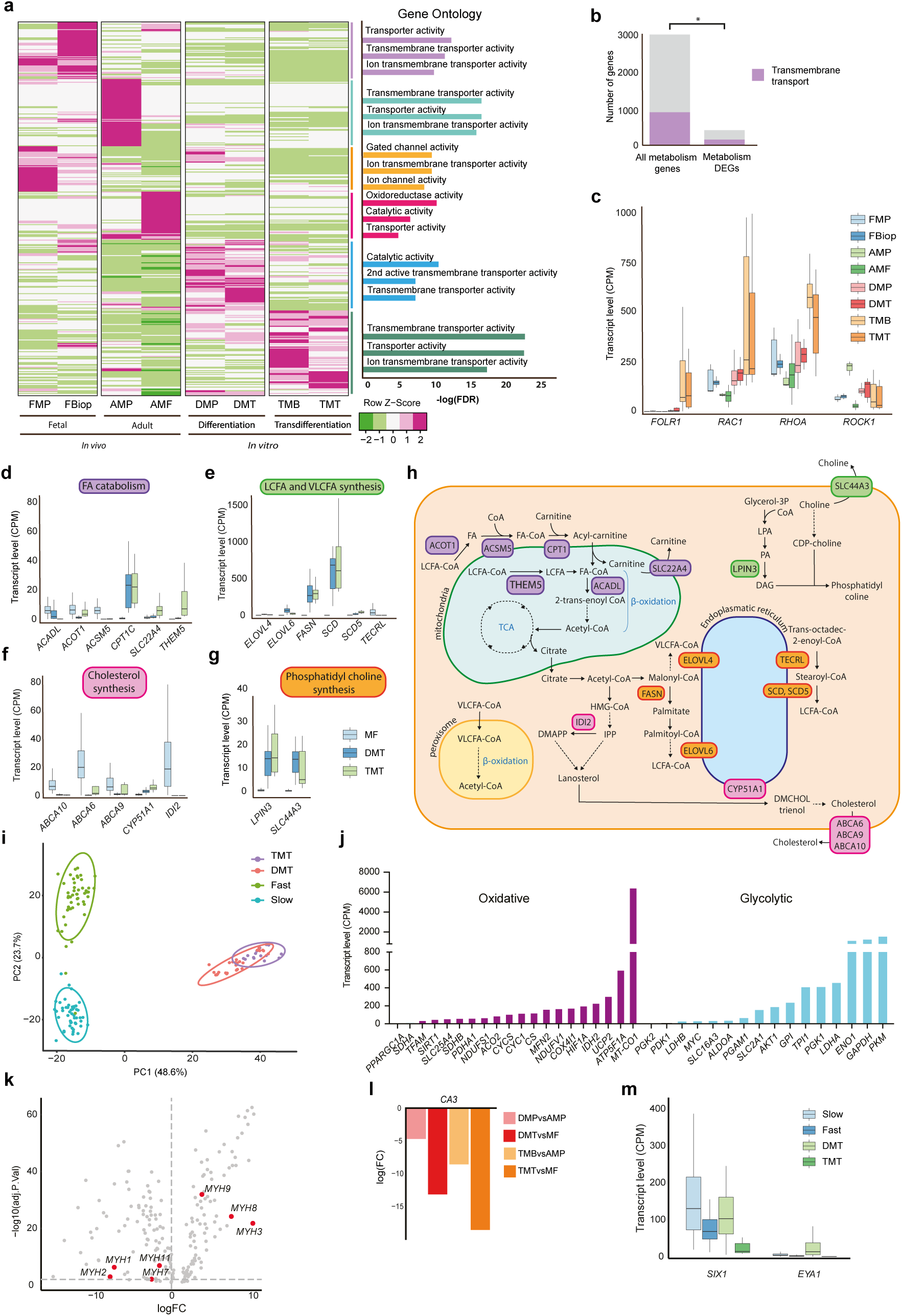
Metabolism and fiber type signatures of in vitro models of human skeletal muscle. **a)** Heatmap showing differentially expressed metabolism genes in the in vitro models, plotted as transformed Z-score for the average CPM across all categories. Differentially expressed gene blocks characteristic for each sample are analyzed individually using Gene Set Enrichment Analysis (GSEA) and the enriched gene ontologies are summarized in the bar plot on the right. **b)** Bar plot highlighting the significant enrichment (*P* = 0.00016) of the proportion of genes associated with transmembrane transport within the differentially expressed metabolism genes as compared to their proportion within all metabolism genes. **c)** Bar plot of average expression levels (CPM) of members of folic acid cycle across in vivo and in vitro samples. **d-g)** Bar pots highlighting average expression levels (CPM) of members of major subprocesses of fatty acid and lipid metabolism, including fatty acid catabolism **(d)**, long chain fatty acid and very-long-chain fatty acid synthesis **(e)**, cholesterol synthesis **(f)** and phosphatidyl choline synthesis **(g)**. **h)** Schematics illustrating lipid and fatty acid cycles and the roles of the genes highlighted in Fig. 3d-g within each respective subprocess. **i)** PCA of isolated human skeletal muscle fiber type 1 and 2 samples compared to transdifferentiated and differentiated in vitro myotubes, based on fiber type-specific marker genes. **j)** Bar plot showing average expression levels (CPM) of genes implicated in glycolytic or oxidative energy metabolism for the hPSC-derived differentiated myotubes. **k)** Volcano plot showing the differentially expressed myogenic genes between the hPSC-derived differentiated myotubes and the adult isolated myofibers, highlighting different Myosin Heavy Chains. **l)** Bar plot demonstrating the logarithmic fold change of *CA3* expression for hPSC-derived differentiated myogenic progenitors compared to adult myogenic progenitors (DMPvsAMP), hPSC-derived differentiated myotubes compared to adult myofibers (DMTvsAMF), fibroblast-derived transdifferentiated myoblasts compared to adult myogenic progenitors (TMBvsAMP) and fibroblast-derived transdifferentiated myotubes compared to adult myofibers (TMTvsAMF). **m)** Bar plot showing the transcript levels (CPM) of *EYA* and *SIX1* genes, for the tissue-isolated fiber types and in vitro differentiated and transdifferentiated myotubes.

Protein homeostasis is another tightly regulated metabolic process in skeletal muscle cells. The Ankyrin repeat and SOCS Box (ASB) gene family encodes subunits of the E3 ubiquitin ligase complex, which has critical functions in protein turnover. Analysis of this gene group revealed that seven members of the ASB family were entirely absent from the in vitro models at both myoblast and myotube stages as compared to either fetal or adult in vivo references, highlighting a major aberration in the protein homeostasis machinery in the in vitro models (Supplementary Figure 3f).

Skeletal muscle fibers can differ in their preferred choice of energy metabolism based on their fiber subtypes. Thus, we aimed to investigate if the in vitro models generated a specific fiber subtype that resembled either oxidative slow-twitch fibers (Type I) or glycolytic fast-twitch fibers (Type II). A principal component analysis based on the genes associated with slow and fast fiber type signatures suggested that hPSC-derived differentiated and fibroblast-derived transdifferentiated myotube cultures differed equally from both slow and fast fiber types and did not differ much from each other for the expression of fiber type-specific genes (Figure 3i)^57–60^. Analysis of the expression of genes related to energy metabolism showed that for both in vitro models, genes related to oxidative and glycolytic metabolism were upregulated, although a bigger fraction of glycolysis-related genes had high expression levels, potentially suggesting that these cultures may be more glycolytic (Figure 3j – Supplementary figure 3h-i).

Myosin heavy chains play an important role in muscle contraction. *MYH1*, *MYH2* and *MYH4* have been identified to be mainly expressed in fast type muscle fibers, whereas *MYH6* and *MYH7* are associated with the slow fiber type^61,62^. Although several of the fast-type related myosin heavy chains were upregulated in the in vitro systems, another group of them were downregulated (Figure 3k and Supplementary figure 3g). However, the expression of *CA3*, a skeletal muscle-specific carbonic anhydrase that is expressed at higher levels in slow fibers, was drastically downregulated in both in vitro models, supporting the notion of a more fast fiber-like phenotype (Figure 3l)^63^.

Finally, we analyzed the expression of the bipartite transcription factor complex *EYA1-SIX1*, which has been shown to promote the reprogramming of slow-twitch fibers to fast-twitch type^64^. We show that the hPSC-derived myotubes have considerable levels of expression of both members of the complex, arguing for a fast fiber type signature (Figure 3m). These analyses suggest that both hPSC-derived differentiation and fibroblast-derived transdifferentiation models recapitulate some aspects of the fast-twitch fiber phenotypes, although their fiber type identity does not strictly adhere to one single type.

### Altered landscape of signaling pathways in models of skeletal muscle

The precise expression of specific signaling pathways is tightly regulated in order to ensure the successful progression of cell fate commitment during development. In the context of developmental and postnatal myogenesis, a handful of signaling pathways such as Wnt, Notch, Sonic Hedgehog and Fibroblast Growth Factor (FGF), have been shown to play indispensable roles in this process^65^. Additionally, supplementation of small molecules to alter or boost signaling pathways during *MYOD1-*induced transdifferentiation and hPSC-derived myogenic differentiation protocols has long been described and has been shown to have beneficial effects on the size and maturation of skeletal muscle cells in vitro^4,66^. To analyze the expression of the members of signaling pathways in the in vitro models, we compiled a comprehensive gene set of 19 pathways, both with well and less known functions in myogenesis. First, we interrogated the different patterns of significantly up- and downregulated members of the individual signaling pathways. We observed that the highest percentage of downregulated pathway members was 40% and included the Insulin Growth Factor (IGF), Notch, Janus kinase/signal transducer and activator of transcription (JAK/STAT), Epidermal Growth Factor (EGF), Fibroblast Growth Factor (FGF), Hepatocyte Growth Factor (HGF), the Toll Like Receptor (TLR) and B cell pathways (Figure 4a). All of these pathways, together with the Hippo pathway, also had up to 60% of their members significantly upregulated in the in vitro models in comparison to the in vivo references (Figure 4b). To reveal the largest discrepancies between the signaling landscape of in vitro models and in vivo references, we utilized stringent expression cutoffs for up- and downregulated signaling pathway members. This analysis showed the Hippo, Notch, FGF, and HGF pathways as the most divergent ones in comparison to the adult myofibers and Notch, JAK/STAT, FGF and Wnt pathways when compared to fetal samples (Supplementary figure 4a-b). Importantly, the adult myofibers differed drastically from all stages of in vitro models, mainly due to their very low expression of the members of these pathways (Figure 4c). In addition, there did not appear to be a common pattern of expression of pathway members between the in vitro models and the in vivo samples, indicating that the changes in the pathways across different comparisons, do not involve the same genes. Although the in vitro samples are known to have a profile which is more akin to that of fetal skeletal muscle, we observed the substantial downregulation of various members of the EGF, FGF and HGF pathways in the transdifferentiated and differentiated myotubes compared to fetal biopsies, suggesting that they do not also fully model fetal myogenic cells at the current state-of-the-art (Figure 4d-e). FGF ligands were most commonly downregulated and therefore, we explored the individual deviations in FGF ligand expression for the different stages of in vitro muscle development compared to fetal myogenic progenitors and full muscle biopsies. Ten FGF ligands were strictly missing in the in vitro models, whereas only one ligand, *FGF5* showed upregulation (Figure 4f). However, the upregulation of *FGF5* was only observed in the transdifferentiated myoblasts compared to fetal myogenic progenitors, while it was not differentially expressed in other comparisons of in vitro models and in vivo references (Supplementary figure 4c). We also observed a group of ligands that were commonly missing in different pair-wise comparisons (Figure 4g). *FGF18,* together with the FGF-activating enzyme *KL,* was the only ligand, which was missing in all stages of in vitro models compared to fetal samples, whereas *FGF13* was only missing in the transdifferentiated myoblasts and *FGF9* in the transdifferentiated myotubes. Furthermore, between late-stage differentiation and transdifferentiation, *FGF4* and *FGF6* were shared as commonly missing ligands. FGF signaling is highly coordinated and ligand-receptor binding is specific. Therefore, the absence of these ligands could exert a limiting effect on myogenesis, as FGF plays a regulatory role in mesoderm fate specification during early development and somite formation^67,68^. Finally, in the differentially expressed genes of the transdifferentiation system compared to both adult and fetal samples, multiple genes related to downstream signaling cascades of receptor tyrosine kinases were identified. These signaling cascades can lead to diverse outcomes, ranging from proliferation and differentiation to survival and cell migration. In transdifferentiated myotubes compared to adult myofibers, two activators of the PI3K/AKT pathway, *SEMA4D* and *PEBP4,* were downregulated, while MAPK/p38 pathway members, *SHC3* and *MAPK11, were* upregulated (Figure 4h). The p38 pathway has been described to promote the *MYOD1* activity, suggesting that the increase in expression of the members of this pathway could be related to MYOD1 activity during MYOD1-induced fibroblast transdifferentiation^69–71^. A comparison between the transdifferentiated cultures and fetal samples also demonstrated the downregulation of the *PLCB2* gene, which is an important activator of the phospholipase C and inositol triphosphate calcium (Ca^2+^) signaling cascade. Ca^2+^ signaling plays a crucial role in the regulation of myoblast differentiation during development but also adult regeneration, suggesting that dysregulation of this pathway might contribute to an incomplete myogenic identity in the transdifferentiation model (Supplementary figure 4d)^72^.

**Figure 4.**
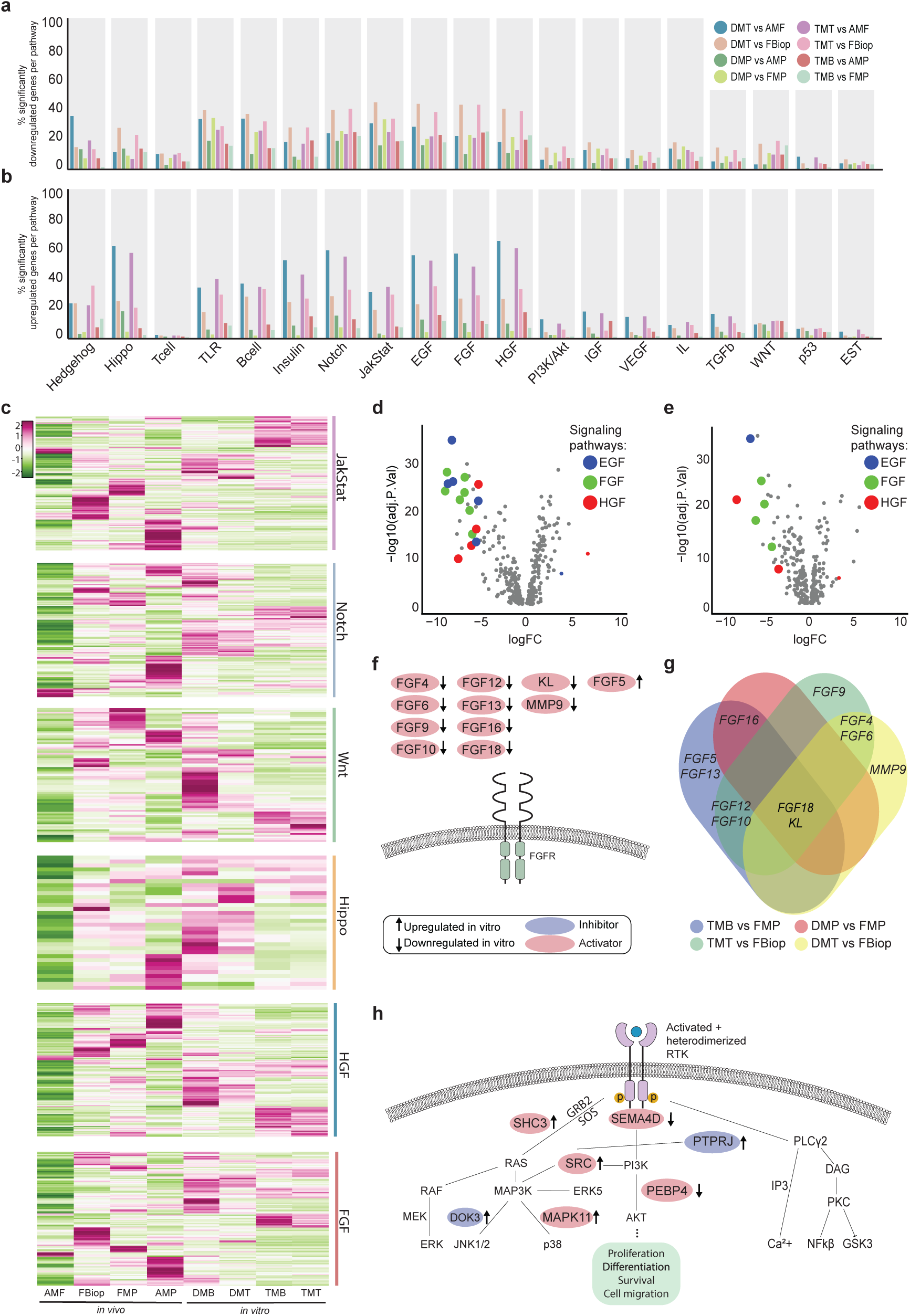
Signaling pathway landscape of in vitro models of human skeletal muscle. **a-b)** Bar plots showing percentages of significantly downregulated **(a)** and upregulated **(b)** members of individual signaling pathways across all indicated comparisons. **c)** Heatmap showing average transcript levels (CPM) of all members of six highlighted signaling pathways in the in vivo and in vitro samples. **d-e)** Volcano plots of differentially expressed genes in transdifferentiated myotubes **(d)** and hPSC-derived differentiated myotubes **(e)** in comparison to fetal biopsies. Significantly downregulated members of the EGF (blue), HGF (red) and FGF (green) signaling pathways are highlighted in color. **f)** Schematics depicting all significantly downregulated (FDR < 0.05) ligands of the FGF signaling pathway for transdifferentiation and hPSC-derived differentiation models at myoblast and myotube stages compared to fetal references. **g)** Venn diagram displaying the individual comparisons that revealed the genes summarized in Fig, 4f, highlighting overlapping FGF ligands across multiple comparisons. **h)** Schematics depicting significantly up- and downregulated members of downstream receptor tyrosine kinase (RTK) signaling for transdifferentiated myotubes as compared to isolated adult myofibers (TMTvsAMF). Red label indicates activators of the pathway, while blue label shows the inhibitors. Up-and downregulated members are indicated by arrows in either direction, respectively.

### Differences in transcriptomic identity of PAX7+ satellite cells in vitro and in vivo

Finally, we aimed to characterize PAX7^+^ populations in the in vitro models in comparison to their in vivo references of human skeletal muscle at different stages of life. In vivo, *PAX7* expression denotes 2 different stages of myogenic progenitor cells. The first one is a group of developmental, proliferating progenitors which are myogenically committed to eventually fuse and form myofibers during embryonic development. The second group consists of the cells that are set-aside, quiescent stem cells, also termed as satellite cells or the resident muscle stem cells, with an important role in adult regeneration. Satellite cells have been proven hard to study in vitro since they quickly lose their quiescent nature after biopsy and isolation procedures^73^. Therefore, research has been dedicated to recapitulating the satellite cell phenotype in vitro to be able to investigate this important cell type. To interrogate the differences of 2D and 3D in vitro primary cell- and hPSC-derived differentiation models in comparison to different stages of human skeletal muscle, we made use of the single cell transcriptomics datasets, which have been instrumental in discerning cell identities within tissues and cellular models. Surprisingly, majority of the limited available single cell transcriptomics datasets are derived from scRNAseq, although the largest fraction of skeletal muscle or its in vitro models is composed of multinucleated myofibers or myotubes, respectively. scRNAseq technology can mainly capture mononucleated cells and as a consequence of this incompatibility, myofiber/myotube-associated transcriptomes are largely dismissed in these analyses, whereas single nucleus RNAseq (snRNAseq) can more faithfully capture the dominant representation of the myofibers within skeletal muscle (Supplementary Figure 1a vs. Supplementary Figure 5a). Based on this comparison, we strongly argue for the use of snRNAseq, when myofiber-associated transcriptomes are studied. Nevertheless, for the interrogation of mononucleated PAX7^+^ skeletal muscle progenitors or stem cells, scRNAseq datasets provide a highly valuable platform. Thus, we integrated six publicly available scRNA sequencing datasets (Figure 5a). In total, 14 clusters were identified across these 6 datasets (Supplementary Figure 5a). All individual clusters were assessed for the percentage of *PAX7* expressing cells and the level of average *PAX7* expression per study and per cluster and eventually three clusters were selected as the most prominent *PAX7*^+^ clusters based on these criteria. Interestingly, *PAX7*^+^ population in Cluster 1 was mainly made up of the cells derived in 3D organoid studies with smaller contributions from the other samples, whilst Cluster 2 was the only cluster with *PAX7*^+^ cells identified in adult human skeletal muscle. Cluster 4, on the other hand, consisted mainly of *PAX7*^+^ cells identified in fetal and embryonic muscle and 2D hPSC-derived differentiation model (Figure 5b). To identify the cell populations in each cluster we performed pairwise differential gene expression analysis between the clusters and additionally looked for the marker genes for each of these three clusters. First, we observed high upregulation of cell cycle-related genes and the proliferative myogenic progenitor marker *ERBB3* in Cluster 4 compared to the other two clusters (Figure 5c)^74^. This proliferative phenotype was supported by the detection of other cell cycle-related genes, such as *CDC7* and *CCNB1,* among the top 20 cluster identifier marker genes (Supplementary figure 5b). Cluster 2, on the other hand, showed the joined upregulation of satellite cell early activation markers, *MYC*, *FOS* and *JUN* and had the highest levels of *MYF5* expression. *MYOG* and *MYOD1* were expressed at low levels in Cluster 1 and 2 and were heavily upregulated in Cluster 4 (Figure 5d and Supplementary Figure 5c). Finally, Cluster 1 was characterized by the differential expression of several extracellular matrix proteins, members of the Notch signaling pathway and markers of satellite cell quiescence, *CXCR4* and *CAV1* (Figure 5e)^75,76^. These observations revealed a continuum of quiescence reaching to a proliferative phenotype, starting from Cluster 1 exhibiting a deep quiescence state, Cluster 2 with a shallow quiescence state and Cluster 4 with a proliferative myogenic phenotype (Figure 5f). Based on these characterizations, we conclude that the *PAX7*^+^ cells in Cluster 4 are mainly proliferating, embryonic myogenic progenitors, while those in Cluster 1 are satellite cells in deep quiescence and those in Cluster 2 are satellite cells at a shallow quiescent state (Figure 5f).

**Figure 5.**
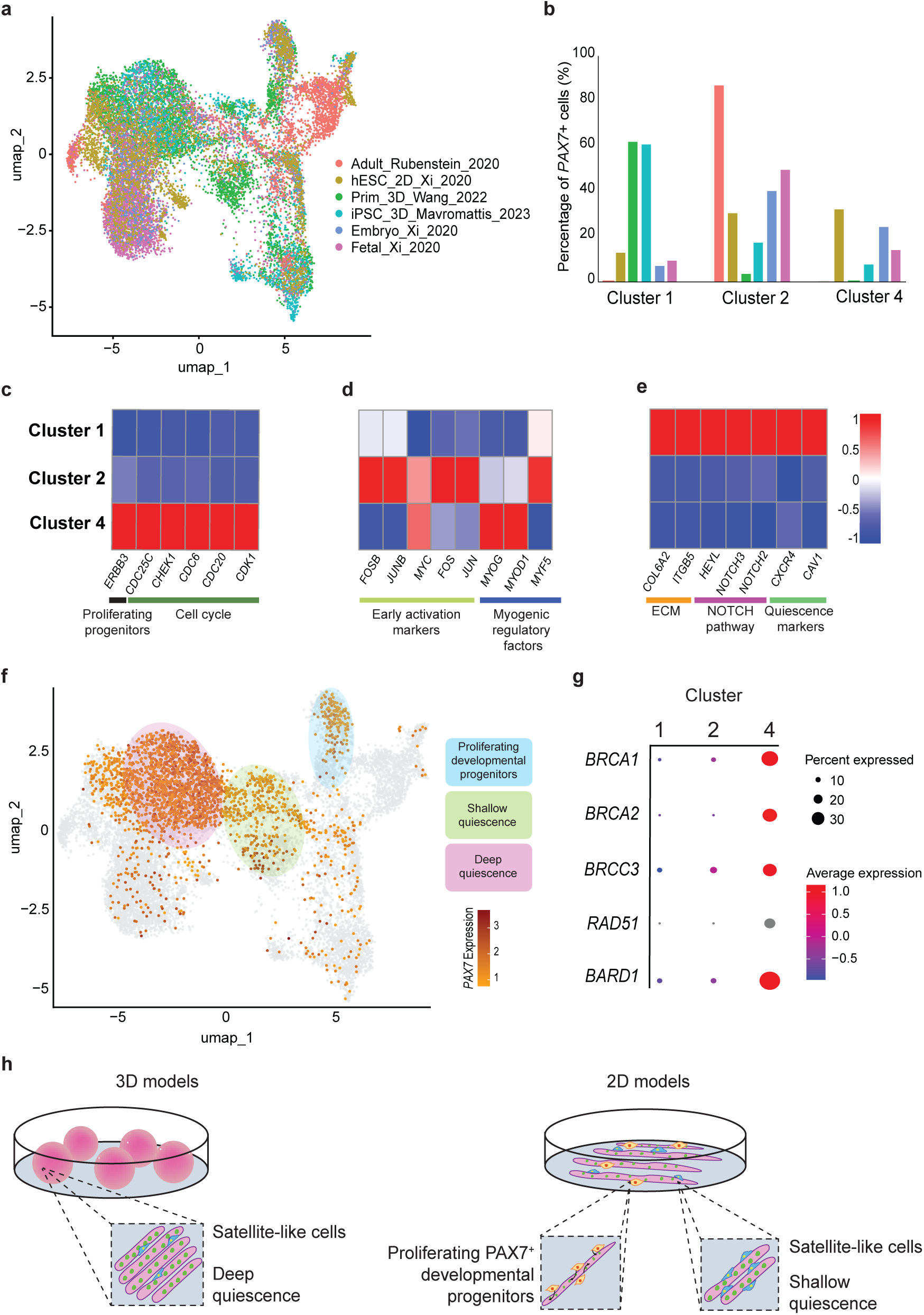
Differences between distinct populations of PAX7^+^ myogenic cells generated in 2D and 3D models of human skeletal muscle. **a)** UMAP showing the integrated scRNAseq studies. The cellular origins of each in vitro model or the developmental stage of in vivo samples are indicated at the beginning of each sample label, followed by the mode of culture model (2D vs. 3D), last name of the first author of the study and the year of publication. (Prim: Primary myogenic cells derived from biopsies). **b)** Bar plot displaying the fraction of PAX7^+^ cells in the indicated clusters within the total number of PAX7^+^ cells in each study. The studies are color-coded as in Fig. 5a. **c-e)** Heatmaps highlighting the differentially expressed genes between the three indicated clusters; genes associated with proliferating developmental progenitors **(c)**, genes related to a shallow quiescent state **(d)** and genes related to a deep quiescent state **(e)**. **f)** Annotation of the clusters of interest: Deep quiescent cluster (1, pink), shallow quiescent and early activated satellite cell cluster (2, green) and proliferating developmental progenitor cluster (4, blue). **g)** Dot plot demonstrating the level and the percentage of expression of genes associated with the BRCA1-BRCA2-containing Complex within indicated clusters. **h)** Model highlighting the generation of satellite-like cells at different quiescent states from 2D and 3D differentiation methodologies.

Surprisingly, we also observed a significant enrichment of expression of five members of the BRCA1-BRCA2-containing complex (BRCC) in Cluster 4, specifically *BRCA1*, *BRCA2*, *BRCC3*, *RAD51* and *BARD1* (Figure 5g). There is limited evidence of the role of BRCC in myogenesis, however, it has been shown that one of the members of BRCC, *BABAM2*, enhanced the differentiation and fusion of adult satellite cells in mouse skeletal muscle regeneration^77–79^. The high expression of BRCC members in Cluster 4, on the other hand, suggests a potential developmental role for this complex during myogenic progenitor commitment and differentiation in human.

Thus, our analyses reveal that the 2D hPSC-derived differentiation model generates a small subset of deep quiescent satellite-like cells and a larger group of shallow quiescent satellite-like cells with similar sized group of proliferating developmental progenitors. Conversely, 3D skeletal muscle organoids, regardless of whether they were derived from hPSCs or primary cells, mainly give rise to satellite-like cells with a deep quiescence signature (Figure 5h).

## Discussion

In vitro models are pivotal to study development, disease and general biological processes. The objective of this study was to discern discrepancies between the current skeletal muscle models and adult, fetal, and embryonic skeletal muscle in human in order to reveal the cellular processes that can be targeted to improve the current in vitro models. We highlighted disparities in different processes such as myogenesis, transcription factors and epigenetics, metabolism, signaling and progenitor identities. To the best of our knowledge, such an extensive meta-analysis of published bulk and scRNAseq datasets derived from several different in vitro models has not been performed to date for human cells. Thus, this study provides novel perspectives on the missing elements in human skeletal muscle models.

Our analyses revealed that on the level of the myogenic identity, early and late-stage myogenic regulatory factors *MYF5* and *MYF6* were missing in the transdifferentiation model. Although the absence of *MYF5* can be explained by the possibility that the transdifferentiation happens without passing through a progenitor state, the lack of *MYF6* expression indicates an incomplete differentiation transcriptome in this model. Developmentally, the expression of *MYF6* comes before that of *MYOD1,* however, in adult myogenesis, *MYF6* expression is regulated by *MYOG* and *MYOD1* expression^80,81^. During transdifferentiation, *MYOD1* and *MYOG* might thus fail to set off *MYF6* expression. It has been previously described that expression of *MYF6* is essential for the expression of a set of genes important for skeletal muscle maturation^82^. Therefore, co-overexpression of *MYF6* and *MYOD1* could be used as a way to enhance transdifferentiation as these two MRFs have different target genes and thus a larger fraction of the muscle specific genes can be activated in the presence of both.

In addition to structural genes found in mature myofibers, both the hPSC-derived differentiation and fibroblast-derived transdifferentiation models surprisingly also lacked a group of genes formerly suggested to be associated with immune and endothelial cells. Our analysis on a snRNAseq dataset revealed that a fraction of these genes was robustly expressed in myofiber nuclei, as well. One such gene is nuclear receptor ROR gamma (*RORC*), which is a transcription factor shown to play an important role in skeletal muscle metabolism by regulating processes related to lipid homeostasis and carbohydrate metabolism^83^. This observation suggests alternative roles for this group of genes within myofibers.

We showed that the in vitro models had aberrant expression of genes related to lipid metabolism and the fatty acid cycle. Lipid metabolism plays an extensive role in skeletal muscle as it is one of the main sources for its energy expenditure in vitro. Aberrant upregulation of genes within several lipid metabolism pathways could be caused by a failure in metabolic reprogramming, in particular in the case of the hPSC-derived differentiation model. Previous evidence suggests that fatty acid synthesis is essential for the survival of hPSCs, whereas skeletal muscle in vivo is dependent on fatty acid uptake from the environment as it lacks the expression of fatty acid synthase (*FASN*)^84,85^. Notably, *FASN* was indeed highly expressed in all in vitro models (Figure 3e, Supplementary Figure 3c). Alternatively, this contrast could be explained by the nutrient availability and the culture media composition. It was shown that hPSC metabolism is dependent on the nutrients that are available and changes in culture conditions could alter the metabolic pathways significantly^86^. Therefore, further optimization of the media composition for in vitro models using hPSC differentiation and fibroblast transdifferentiation, but also for differentiation of immortalized muscle cell lines, could enhance their metabolic similarities to bona fide muscle.

The upregulation of folic acid receptor 1 (*FOLR1*) in the in vitro models compared to in vivo adult references could stem from the lack of folate in the culture media. Indeed, folic acid has been shown to be important for the proliferation and differentiation of in vitro cultures^87^, arguing for the supplementation of folate to the culture media as a potential improvement.

The ASB genes were consistently identified as missing in the in vitro models compared to in vivo references. The ASB gene family encodes subunits of the E3 ubiquitin ligase complex and most of its family members have been shown to have high expression in muscle^88^. One of the ASB genes, *ASB15* has been shown to promote muscle differentiation by regulating protein turnover^89^.

Our analyses revealed that 8 FGF ligands were completely missing (Figure 4f) in the in vitro models and highlighted the subtype per in vitro model (Figure 4g). The role of FGF signaling in myogenesis has been rather controversial. FGF has been shown to both positively and negatively affect muscle differentiation in multiple model organisms and in in vitro culture^90–92^. Additionally, FGF2 is already an important component of hPSC-derived muscle differentiation protocols^93–95^. Since the individual ligands likely have different effects, the addition and incubation time of FGF ligands in the culture media should be extensively tested.

Cell fate changes are accompanied by major alterations in the epigenetic landscape. The epigenetic memory of the source cells can be retained in experimental models, and in the case of fibroblast transdifferentiation towards the myogenic lineage, this has been shown to have an inhibitory effect^29^. Thus, we investigated the transcription factors and epigenetic complexes that are aberrantly expressed in the in vitro models. Two major transcription factor families were identified, the *HOX* and *ANKRD* family. Interestingly, a similar analysis which highlighted the differences between in vitro myotubes and muscle biopsies from 5 different patients also pointed to the downregulated expression of *ANKRD* genes^96^. One member of this family is muscle-specific and was found to be downregulated in hPSC-derived differentiation model, while for the other differentially expressed members of the family no clear function in muscle has been described so far. *HOX* genes are tightly regulated during muscle development. They are expressed in a temporo-spatial manner and are important for tissue patterning. Their roles during human myogenesis are less well understood, although in *D. melanogaster* it was shown that expression of one of the *HOX* genes, Ultrabithorax (*Ubx*), was essential for setting off the myogenic differentiation program by downregulating main regulator, *twist*, in the adult progenitors^97^. Our analysis showed differential expression for 13 individual HOX genes, across all four individual clusters *HOXA*-*D,* indicating that they do have a role during human myogenesis, and that potential temporal dysregulation of these genes might affect the efficiency of in vitro muscle differentiation.

HOX genes are regulated in part by Polycomb Repressive Complex 1 (PRC1)^98^. We showed that canonical PRC1 components (*BMI1*, *CBX6*, *PCGF2*, *PHC2*, *RING1* and *SCMH1*) were upregulated in the in vitro models and this high expression was already present in their source cells, namely the hPSCs and fibroblasts, and failed to be erased upon differentiation or transdifferentiation, respectively. PRC1 stabilizes the expression of cell fate commitment genes. Therefore, chemical inhibition of the PRC1 complex could potentially enhance transdifferentiation by allowing the activation of muscle differentiation-specific gene networks. Another major epigenetic complex which plays an important role in muscle differentiation is the SWI/SNF ATP dependent chromatin remodeling complex^99^. Its two main facilitators in myogenesis are *SMARCA2* and *SMARCA4.* These two SWI/SNF enzymes have distinct functions in myogenesis and are indispensable for the myogenic program*. SMARCA4* activates muscle gene transcription at the earlier stages of myogenesis whilst *SMARCA2* causes proliferating myoblasts to exit the cell cycle by repression of cyclin D1 (*CCND1*). Specifically, *SMARCA2* expression was absent in all stages of differentiation and transdifferentiation, suggesting that the cells may not be fully exiting the cell cycle, which in turn would limit their fusion capacity.

Finally, we analyzed an integrated dataset of six different scRNAseq samples, covering 4 studies to identify transcriptomic identities of PAX7^+^ cells in the in vitro models. In the cluster containing the proliferating developmental PAX7^+^ progenitors, we also identified the differential expression of the BRCA1-BRCA2-containing complex (BRCC), which encompasses *BRCA1* and *BRCA2* and *BRCC3* in addition to the cell cycle genes. The BRCC complex has been described to play a role in skeletal muscle metabolism ^78,79^. Due to its high expression in the differentiating cluster, we hypothesize that this complex likely plays an important role during the myogenic commitment in embryogenesis.

Interestingly, we observed that the 3D organoid models mainly generated satellite-like cells with a deep quiescence phenotype, marked by the expression of the NOTCH pathway and extracellular matrix genes^100–102^. Conversely, hPSC-derived 2D differentiation model generated 1) a population of satellite-like cells at a shallow quiescence state, clustering together with the adult PAX7^+^ cells, 2) proliferating developmental PAX7^+^ progenitors, clustering together with the majority of the fetal and embryonic PAX7^+^ cells and finally also 3) a very small group of satellite-like cells with a deep quiescence phenotype. Therefore, we argue that 2D cultures give rise to a diverse PAX7^+^ cell population, while 3D cultures mainly produce PAX7^+^ satellite-like cells that show a deeper quiescence profile, potentially due to the enhanced extracellular matrix niche within the model.

Surprisingly, although a similar approach of cell isolation was used to dissociate the cells from the adult muscle and the 3D organoids, the adult muscle samples did not include any cells which showed a deep quiescent profile. We speculated that this might be due to the initial insult of the biopsy process, which might set off an early activation response of satellite cells in vivo. The accumulation of more adult scRNAseq studies could reveal a PAX7^+^ population in adult muscle, comprised of deeply quiescent satellite cells.

In conclusion, this large-scale meta-analysis covering more than 400 published bulk RNAseq and seven scRNAseq or snRNAseq datasets, uncovered differences between in vitro human skeletal muscle models and in vivo references. Systematic characterization of these differences provides novel insights regarding skeletal muscle cell identity, while also suggesting adjustments in the protocols to improve the existing models. This holistic approach can also be used for other somatic cells in human and their in vitro models, paving the way for a better understanding of cellular identities and existing models.

## Methods

### Curation of the Dataset

To perform the large-scale analysis of in vitro models of human skeletal muscle in comparison to in vivo samples, a thorough literature search led to a collection of a total of 418 bulk RNA sequencing samples covering 34 independent studies^12–31,33,34,36–46,103^. The sample types included in our sample collection were fibroblasts, adult, fetal and embryonic skeletal muscle biopsies of complete tissue and their sorted myogenic cells, human pluripotent stem cells, primary cultures of myogenic cells, immortalized myogenic cell lines and myogenic cultures derived from differentiation and transdifferentiation protocols for human cells of different stages. For the in vivo references, only healthy control samples were included. These samples were downloaded as raw FASTQ files from the Gene Expression Omnibus database (GEO; https://www.ncbi.nlm.nih.gov/geo/) using the GNU command line tool Wget and samples were labelled according to the study and cell type to facilitate identification. Additionally, for the integration and analysis of six scRNA sequencing datasets, scRNA-seq count matrices and associated metadata were downloaded from the sources, which are provided in the Data Availability section. Finally, one single nucleus RNA (snRNA) sequencing study was used to confirm the expression of predicted immune system and endothelial cell-related genes within the myofiber-associated cluster. FastQC (package version 0.12.0). was used for the quality control of all bulk RNAseq data^104^ .

### Preprocessing of Data

A count table was generated by aligning the samples to the human reference genome (Gencode, release 44, (GRCh38.p14)) using STAR aligner (version 2.5.26). Next, featureCounts program (version 2.0.1) was used to count the number of aligned reads to the reference genome. All samples were mapped using a principal component analysis (PCA) after normalization and logarithmic transformation. Individual pairwise comparisons were selected from the master count table and raw gene counts, later converted to normalized counts per million (cpm), were used as input for further analysis. The pairwise comparisons we performed include: 1) “myogenic progenitors from hPSC differentiation vs. fetal and embryonic muscle samples”, 2) “myotubes from hPSC differentiation vs. sorted myofibers from biopsies”, 3) “myotubes from hPSC differentiation vs. skeletal muscle biopsies”, 4) “myoblasts from fibroblast transdifferentiation vs. fetal and embryonic muscle samples”, 5) “myotubes from fibroblast transdifferentiation vs. sorted myofibers from biopsies”, 6) “myotubes from fibroblast transdifferentiation vs. skeletal muscle biopsies”, 7) “immortalized cell line myoblasts vs. primary myoblasts”, 8) “immortalized cell line myoblasts vs. fetal and embryonic muscle samples”, 9) “immortalized cell line myotubes vs. skeletal muscle biopsies” and 10) “immortalized cell line myotubes vs. sorted myofibers from biopsies”. Additionally, expression profiles from source cells for the in vitro generation methods (fibroblasts and human pluripotent stem cells) were included as an initial point of comparison.

### Differential Gene Expression Analysis and Gene Set Curation

To identify the differentially expressed genes between the in vitro skeletal muscle models and their in vivo counterparts, the integrated Limma^105^, version 3.5.1) and EdgeR^106^, version 4.0.16) workflow was used^107^. Only genes with an adjusted p-value (FDR) smaller than 0.05 were used for further analysis. The significantly differentially expressed genes were filtered for their gene expression levels using counts per million (CPM) filters. Downregulated genes had a median CPM expression of <1 for all samples of the group and therefore were virtually not expressed, whilst for the upregulated genes, a minimum median expression of CPM > 2 was applied for all samples of the group. This resulted in a list of differentially expressed genes (DEGs) (up- and downregulated) per comparison. To identify functions and families of DEGs, lists were filtered for gene sets. Curated gene sets used included, myogenic genes^108,109^, metabolic genes^110^, transcription factors^111^ and epigenetic complexes^112^. In addition, we compiled gene sets for a comprehensive list of signaling pathways, including Sonic Hedgehog, Hippo, PI3K-Akt, T cell receptor, Toll-like receptor, B cell receptor, Insulin, Notch, JAK/STAT, EGF, IGF, VEGF, HGF, FGF, Akt-MTOR, Interleukin, TGFβ, WNT, P53, and the estrogen signaling pathways^113^.

### Downstream Analysis of the Differentially Expressed Genes

Gene ontology analysis was performed using the online Gene Set Enrichment Analysis tool (GSEA; https://www.gsea-msigdb.org/gsea/) and the accompanying R tools MsigdbR^114^, clusterProfiler^115^ and fgsea^116^. In addition, DEG lists were analyzed using the online tool STRING (Szklarczyk et al., 2019, version 12.0, https://string-db.org/) to visualize the predicted protein-protein interaction networks within the up- and down-regulated DEGs.

### scRNAseq: Preprocessing of Datasets

The single-cell RNA-Seq analysis was conducted using the Seurat package (version 5.0.2), beginning with the preprocessing of six individual datasets^36,46–48^. First, three files were used as an input for each dataset: the expression matrix, feature annotation, and cell metadata. Quality control was then performed to filter out low-quality cells, specifically those with fewer than 200 detected features, more than 2,500 detected features, or more than 5% mitochondrial gene content. Normalization of gene expression was done using the LogNormalize method, which scaled the data by the total expression in each cell, multiplied by a scale factor of 10^4^ and transformed it to a log scale. Highly variable features were identified using the variance-stabilizing transformation method (FindVariableFeatures) with 2,000 top genes selected for downstream analysis. The data was then scaled to center and standardize the expression values, and dimensionality reduction was performed using PCA. The first 10 principal components were selected based on their variance contribution, and these were used to construct a shared nearest neighbor (SNN) graph with FindNeighbors. Clustering of cells was performed using the Louvain algorithm (FindClusters) at a resolution of 0.5, and clusters were visualized using Uniform Manifold Approximation and Projection (UMAP) generated with RunUMAP. Marker genes specific to each cluster were identified with FindAllMarkers, requiring a minimum log-fold change of 0.25 and expression in at least 25% of cells within a cluster. The results were visualized using feature plots. Each dataset was saved as a Seurat object in RDS format after preprocessing for subsequent integration.

### scRNAseq: Dataset Integration

To integrate the six datasets into a single Seurat object, the preprocessed RDS files were loaded, and anchors were identified using the FindIntegrationAnchors function, aligning shared cell states across datasets. Batch effects and technical variations were corrected using the IntegrateData function, generating a unified integrated assay. The integrated dataset was scaled and reanalyzed using PCA. A SNN graph was constructed using the integrated data, and clustering was performed again with the Louvain algorithm at a resolution of 0.5, producing integrated clusters. UMAP was applied to visualize the clusters in two dimensions, and additional UMAP plots split by study were generated to evaluate dataset-specific contributions to the integrated clusters. Marker genes for each integrated cluster were identified using the same criteria as in the individual analyses, and their expression was visualized using feature plots. Cluster composition analysis was performed to assess the proportion of cells from each dataset within the integrated clusters, and the results were visualized using bar plots. Specific clusters of interest were extracted for further analyses, including differential expression analysis between clusters and detailed visualization of gene expression patterns. To determine the clusters that are highly expressing the *PAX7* gene, UMAP was applied to visualize the cells expressing *PAX7* in the clusters with a threshold of LogFC > 0.75.

### scRNAseq: Differential Gene Expression Analysis

To identify genes that are differentially expressed between cells in two distinct clusters with the aim of assessing potential phenotypic differences between these groups, first, the gene of interest, *PAX7*, was defined and the layers of data were mapped to their corresponding studies. The count matrix for each layer was extracted using LayerData from the integrated dataset and overlapping cells between the layer and study-specific metadata were identified. If overlapping cells were found, counts were combined across layers. CPM values for *PAX7* were calculated by normalizing raw counts by the library size (total counts per cell) and multiplying by 10^6^. Metadata for each study was updated with the CPM values of *PAX7*, and the processed metadata was added back to the integrated Seurat object using AddMetaData. *PAX7^+^* cells were identified by selecting cells with *PAX7* expression greater than zero using the WhichCells function. These cells were further subset to clusters of interest using “subset” function, ensuring that the default assay was set to “RNA”. Layers in the subset were combined using JoinLayers, and the data was normalized with the LogNormalize method and a scale factor of 10^4^. For cluster-specific analyses, *PAX7^+^* cells from cluster 2 in the “Adult_Rubenstein, 2020” study were identified based on metadata and combined with all *PAX7^+^* cells in clusters 1 and 4. The subsets were merged into a single Seurat object for comparative analysis. To ensure compatibility for downstream analysis, the RNA assay was set as the default, and layers were joined before re-normalizing the data. Differential expression analysis was conducted using the FindMarkers function, applying a minimum log fold-change threshold of 0.25 and requiring a minimum of 25% of cells to express the gene to identify genes differentially expressed between clusters.

### Statistics and reproducibility

For bulk and single cell RNAseq datasets, DEG analyses were performed using EdgeR and Seurat respectively. Differential expression was considered significant for false discovery rate (FDR)-corrected *P* values less than 0.05. Figures 1c, 2c-d, S2b-d, 3c-g, l and S3a, c-f contain box plots that demonstrate the average counts per million (CPM) expression per group and the standard error for the mean (SEM). Pairs of sample groups were compared to each other with a Student’s *t*-test, and the differences were considered significant for *P* values less than 0.05. In Figure 3b, Proportion test was used to assess the significance of percentages.

### Data availability

All datasets used in this study are publicly available and can be downloaded from GEO or BioProject with the following accession codes: GSE129505, GSE121154, GSE111163, GSE87365, GSE161025, GSE234616, GSE221912, GSE178784, GSE93263, PRJNA610985, GSE158216, GSE214495, GSE236120, GSM1527072, GSE128844, GSE98509, GSE86356, GSE130646, GSE78158, GSE78644, GSE102812, GSE78649, GSE89588, GSE100943, GSE112101, GSE117609, GSE117382, GSE114938, GSE163213, GSE119402, GSE136807, GSE124072, GSE235781, GSE18927, GSE147513, GSE147514, GSE188215, GSE147457, GSE130646). SRA numbers of individual samples can be found in Supplementary Table 1 with their respective GEO accession number.

## Acknowledgements

We thank A. Yildirim and M. Di Gloria for their assistance with data organization and gene set generation and B. van der Veer for his assistance with setting up the computing environment. Funding: This work was supported by the Research Foundation – Flanders (FWO, Fonds voor Wetenschappelijk Onderzoek – Vlaanderen, G0DCO23N). A.M.A. is supported by the FWO Doctoral Fellowship. A.Y. is Collen-Francqui Docent.

## Author contributions

M.V.P., A.M.A., E.E.M., J.B and A.Y. designed the analyses. M.V.P., A.M.A., E.E.M. and J.B. curated and analyzed the data. M.V.P., A.M.A., E.E.M., J.B., S.J. and

A.Y. interpreted the data. M.V.P. and A.Y. wrote the manuscript with input from all authors. A.Y. supervised the study. Competing interests: The authors declare no competing interests.

## Notes

### Competing Interest Statement

The authors have declared no competing interest.

