## Supplementary Figures for "Delineating transcriptomic signatures of in vitro human skeletal muscle models in comparison to in vivo references"

**Supplemental information**

**Figure S1: Analysis of single nucleus RNA sequencing data of adult skeletal muscle for the expression of putative myofiber-associated genes a)** UMAP of single nucleus RNA sequencing of adult skeletal muscle sample (Pass, C. G., et al. 2023). Cell identities in clusters are determined by the expression of the marker genes based on the original study and clusters are color-coded for their unique cell types. **b-k)** UMAPs highlighting genes from the predicted immune and endothelial system-related gene lists in Fig. 1e, which show high expression in the myofiber cluster. **l)** Collection of violin plots showing Z-score transformed transcript levels highlighting the genes suggested to be related to immune and endothelial systems in Fig. 1e, but show moderate expression in the myofiber cluster. The presence of a vertical line indicates detected gene expression in a given cluster.

**Figure S2: Analysis of transcription and epigenetic factors in immortalized myogenic cell lines, hPSC- or transdifferentiation-derived myogenic cultures and their in vivo references. a)** Dot plot showing the positive and negative standard logarithmic fold change of gene expression for differentially expressed HOX genes in immortalized cell lines in comparison to adult and fetal in vivo references. **b-d)** Bar plot showing expression levels of individual active members of SWI/SNF **(b)**, PRC1 **(c)** and COMPASS/MLL **(d)** respectively, for all different categories in vivo and in vitro.

**Figure S3: Metabolic and fiber type signatures in immortalized myogenic cell lines, hPSC-derived differentiated and fibroblast-derived transdifferentiated myogenic cultures. a)** Bar plot showing average transcript levels (CPM) of folic acid cycle members for adult myofibers and myotubes derived from immortalized cell lines. **b-e)** Bar plots displaying average transcript levels (CPM) of the members of fatty acid and lipid metabolism subprocesses for adult myofibers and myotubes derived from immortalized cell lines: fatty acid catabolism **(b)**, long chain fatty acid and very-long-chain fatty acid synthesis **(c)**, cholesterol synthesis **(d)** and phosphatidyl choline synthesis **(e)**. **f)** Dot plot showing the standard logarithmic fold change of expression of ASB family of E3 ubiquitin ligases across the indicated comparisons. **g)** Volcano plot showing the differentially expressed myogenic genes between transdifferentiated myotubes and the adult isolated myofibers, highlighting different Myosin Heavy Chains. **h-i)** Bar plot showing average expression levels (CPM) of genes implicated in glycolytic or oxidative energy metabolism for transdifferentiated myotubes.

**Figure S4: Analysis of aberrant expression of signaling pathway members in the in vitro models. a-b)** Bar plots showing the number of significantly up- and downregulated members of each signaling pathway, which passed strict median CPM filtering (>5 CPM in the upregulated fraction, <1 CPM in the downregulated fraction) for in vitro comparisons to adult **(a)** and fetal **(b)** references. **c)** Individual schematics showcasing the differentially expressed ligands of the FGF pathway for transdifferentiated myoblasts compared to fetal myogenic progenitors (upper left), hPSC-derived differentiated myogenic progenitors compared to fetal myogenic progenitors (upper right), transdifferentiated myotubes compared to fetal biopsies (lower left) and hPSC-derived differentiated myotubes compared to fetal biopsies (lower right). **d)** Schematics highlighting the differentially expressed members of the downstream RTK signaling cascades for transdifferentiated myoblasts compared to fetal myogenic progenitors (left) and transdifferentiated myotubes compared to fetal biopsies (right).

**Figure S5: Integration of in vitro and in vivo scRNAseq datasets and characterization of the PAX7$^{+}$ clusters. a)** UMAP showing 14 clusters identified when 6 scRNAseq datasets of in vitro models and in vivo references are integrated. **b)** Dot plot showing the top 20 marker genes of each cluster, average expression is denoted by color and the percentage of gene expression in the cluster is displayed by the dot size. **c)** Bar plot demonstrating the average expression of *MYOD1* for the *PAX7*$^{+}$ cells in each cluster per study.

**Figure S1: Analysis of single nucleus RNA sequencing data of adult skeletal muscle for the expression of putative myofiber-associated genes**

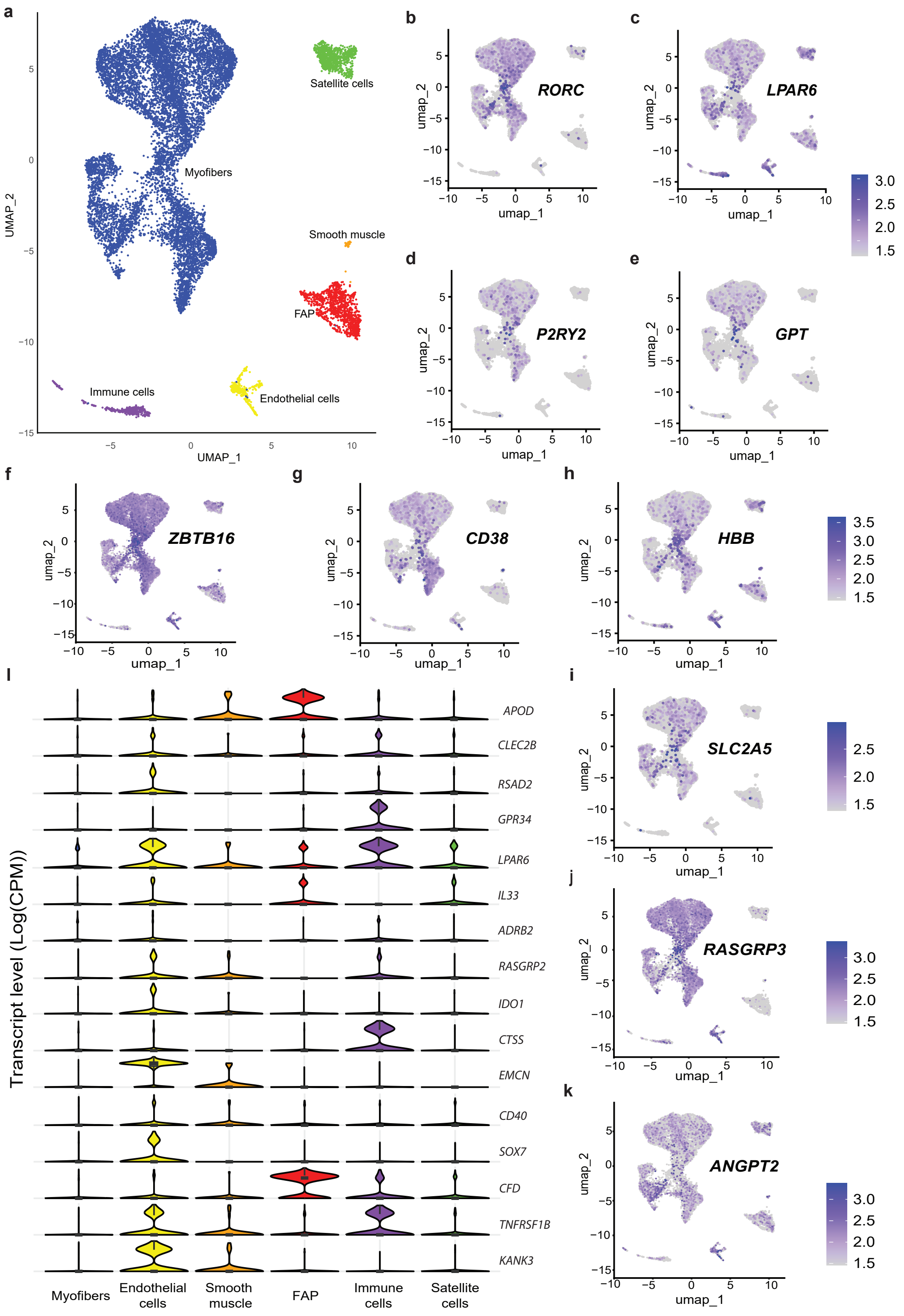

**Figure S2: Analysis of transcription and epigenetic factors in immortalized myogenic cell lines, hPSC- or transdifferentiation-derived myogenic cultures and their in vivo references.**

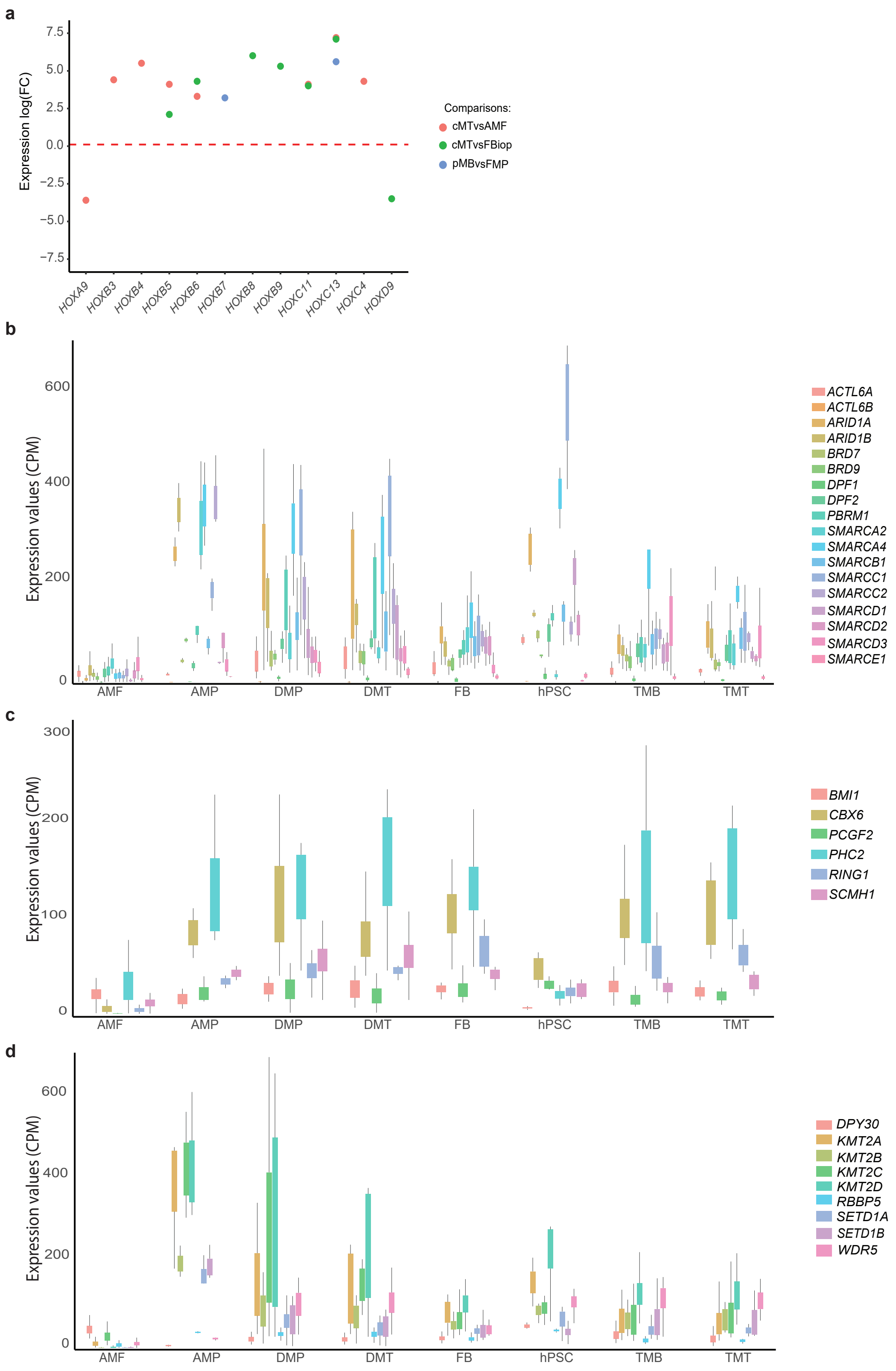

**Figure S3: Metabolic and fiber type signatures in immortalized myogenic cell lines, hPSC-derived differentiated and fibroblast-derived transdifferentiated myogenic cultures.**

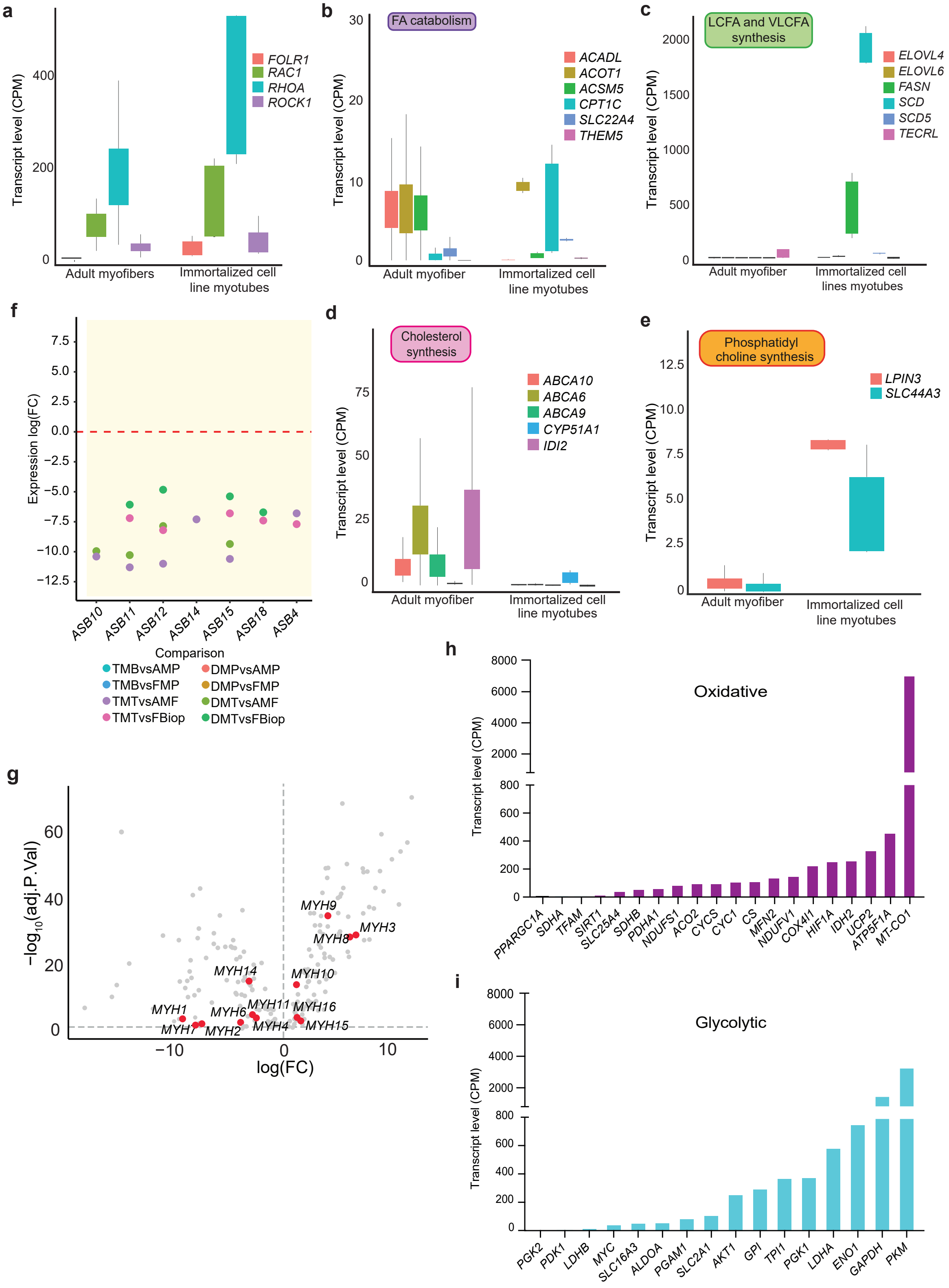

**Figure S4: Analysis of aberrant expression of signaling pathway members in the in vitro models.**

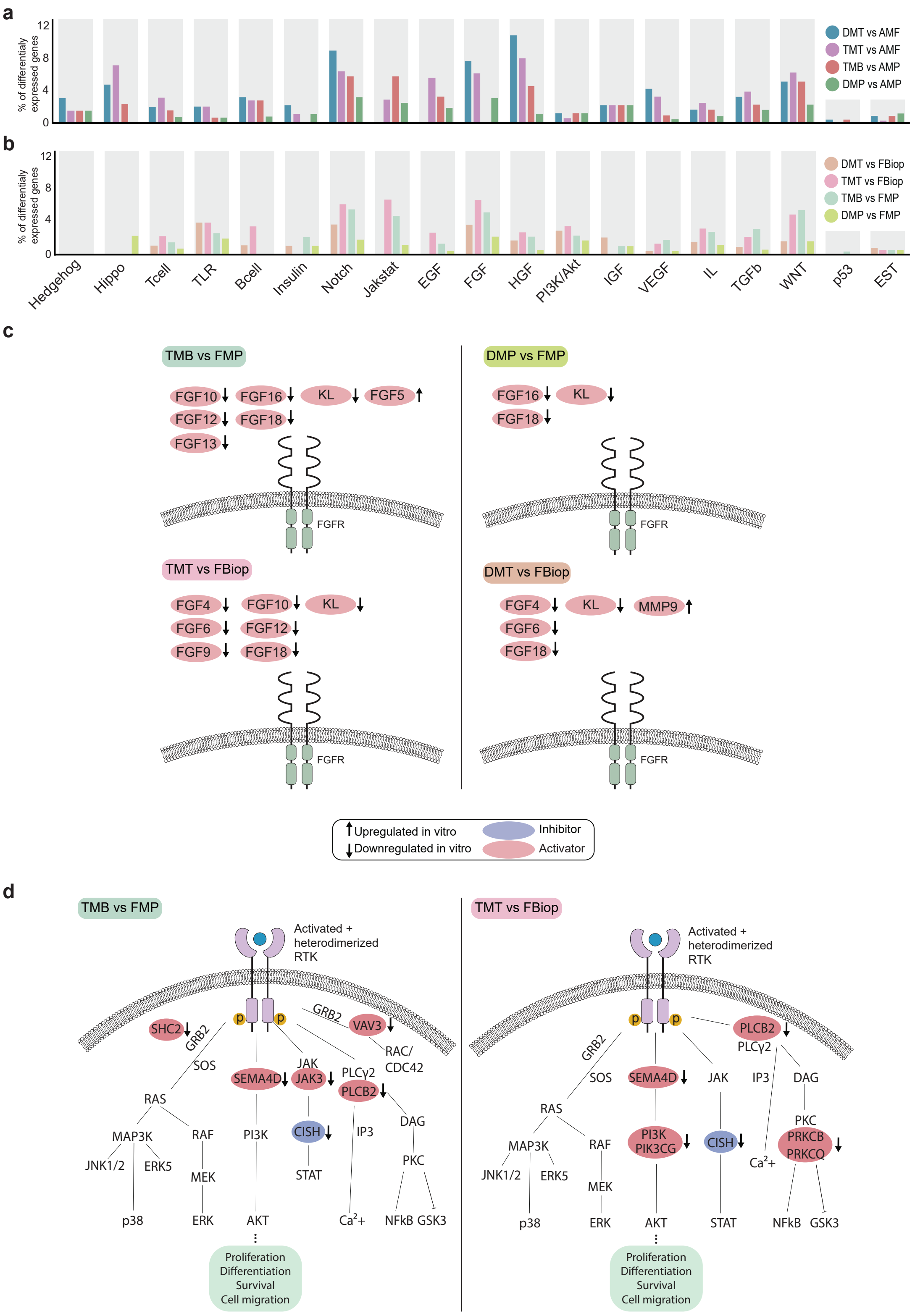

**Figure S5: Integration of in vitro and in vivo scRNAseq datasets and characterization of the PAX7$^{+}$ clusters.**

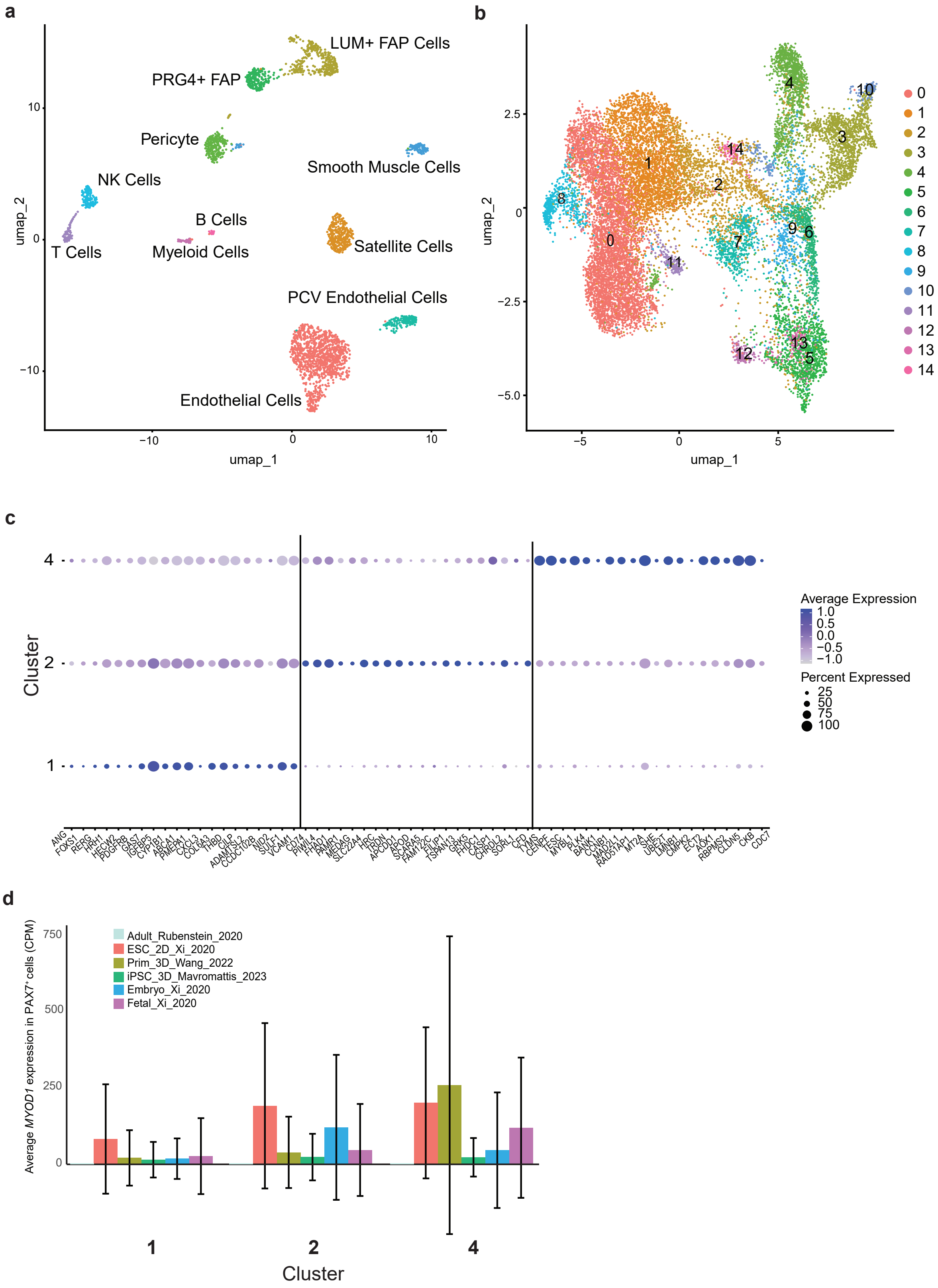
